# Incursion pathways of the Asian tiger mosquito (*Aedes albopictus*) into Australia contrast sharply with those of the yellow fever mosquito (*Aedes aegypti*)

**DOI:** 10.1101/2020.03.27.012666

**Authors:** Thomas L Schmidt, Jessica Chung, Anthony R. van Rooyen, Angus Sly, Andrew R Weeks, Ary A Hoffmann

## Abstract

**BACKGROUND:** Understanding pest incursion pathways is critical for preventing new invasions and for stopping the transfer of alleles that reduce the efficacy of local control methods. The mosquitoes *Aedes albopictus* (Skuse) and *Ae. aegypti* (Linnaeus) are both highly invasive disease vectors, and through a series of ongoing international incursions are continuing to colonise new regions and spread insecticide resistance alleles among established populations. This study uses high-resolution molecular markers and a set of 241 reference genotypes to trace incursion pathways of *Ae. albopictus* into mainland Australia, where no successful invasions have yet been observed. We contrast these results with incursion pathways of *Ae. aegypti*, investigated previously.

**RESULTS:** Assignments successful identified China, Japan, Singapore and Taiwan as source locations. Incursion pathways of *Ae. albopictus* were entirely different to those of *Ae. aegypti*, despite broad sympatry of these species throughout the Indo-Pacific region. Incursions of *Ae. albopictus* appeared to have come predominantly along marine routes from key trading locations, while *Ae. aegypti* was mostly linked to aerial routes from tourism hotspots.

**CONCLUSION:** These results demonstrate how genomics can help decipher otherwise cryptic incursion pathways. The inclusion of reference genotypes from the Americas may help resolve some unsuccessful assignments. While many congeneric taxa will share common incursion pathways, this study highlights that this is not always the case, and incursion pathways of important taxa should be specifically investigated. Species differences in aerial and marine incursion rates may reflect the efficacy of ongoing control programs such as aircraft disinsection.

## 1 Introduction

Incursions of invasive pests present a range of threats to local biodiversity, agriculture and human health.^1–4^ Pest incursions into new regions threaten the colonisation of these regions, while incursions into regions with established populations threaten the introduction of advantageous alleles that reduce the efficacy of control programs.^5^ Following these criteria, an incursion may not only refer to the dispersal of exotic taxa but to the dispersal of any taxa with consequences for pest management. Dispersing pests are frequently transported as “stowaways” on trade and transport vessels,^6–8^ and the distributions of these species have increased accordingly with the upsurge in recent decades of trade and transport activity.^9^ Intercepting incursive material at borders can be an effective means of stopping incursions,^10^ but is resource-intensive. Knowledge of incursion pathways is vital for strategic deployment of these resources, allowing inspections to focus on likely source locations^11^ and common entry points.^12^

The Asian tiger mosquito (*Aedes albopictus*, Skuse) is a hyperaggressive dengue vector that has successfully invaded tropical and temperate regions worldwide from its native range in Asia.^13^ It is considered one of the world’s most dangerous invasive species (Global Invasive Species Database, http://www.issg.org/database/). In Australia, border security makes frequent interceptions of *Ae. albopictus* at international ports across the country.^14^ While much of the Australian mainland likely presents suitable habitat for *Ae. albopictus,*^15^ no successful establishment has yet been recorded. However, an invasion of *Ae. albopictus* was recently recorded in the Torres Strait Islands,^16^ where its present distribution is only tens of kilometres from the Australian mainland.^14^

A recent study that traced incursion routes of the congeneric mosquito *Ae. aegypti* (Linnaeus) into Australia found that most (62) had come from Bali, Indonesia, with others (21) mostly assigned to other locations in Southeast Asia.^8^ These assignments were derived using a reference panel of *Ae. aegypti* genotypes taken predominantly from locations with direct flights into Australia. Certain locations with direct flights and presumably large *Ae. aegypti* populations, such as Singapore, Taiwan, and Ho Chi Minh City, were not found to be source populations of any of the *Ae. aegypti*.

Although *Ae. albopictus* and *Ae. aegypti* both disperse internationally on vessels like ships and aeroplanes,^17–19^ analysis of their genetic structure in the Indo-Pacific has indicated that gene flow pathways are different in these species.^20^ Likewise, interceptions at New Zealand borders found that most *Ae. albopictus* were transported as larvae on ships, while most *Ae. aegypti* were transported as adults on aeroplanes.^17^ One potential reason for these differences is that *Ae. albopictus* may be able to withstand longer transportation times and colder minimum temperatures by undergoing diapause,^21–23^ an option unavailable to *Ae. aegypti.*^24^ Diapause has also been proposed as a means by which *Ae. albopictus* has been able to colonise distant locations without undergoing ‘stepping stone’ colonisation of intermediate sites.^20,25–27^ If gene flow proceeds along different pathways for *Ae. albopictus* and *Ae. aegypti*, incursion routes may also be different, even if both species are locally abundant and can be transported by the same vessels.

This study investigates *Ae. albopictus* incursion pathways into mainland Australia, and contrasts these with previously determined incursion pathways of *Ae. aegypti* (c.f. Schmidt et al.^8^). By genotyping *Ae. albopictus* detected at Australian borders, and comparing these genotypes with a panel of reference genotypes from across the Indo-Pacific region, the likely source population of each mosquito can be identified. Accurate identification is conditional on several factors, including sufficiently high-quality DNA of the intercepted sample, a wide geographic range of reference genotypes, and sufficient genetic differentiation among the reference genotypes. This latter factor is important to consider in assigning *Ae. albopictus*, as *Ae. albopictus* populations are typically less differentiated than those of *Ae. aegypti*,^20,28^ which may lead to lower precision when assigning *Ae. albopictus*. Nevertheless, the strong differentiation between Indonesian and non-Indonesian populations of *Ae. albopictus*^20^ should allow for assignment between these groups, and thus determine whether *Ae. albopictus*, like *Ae. aegypti*, are transported to Australia predominantly from nearby Indonesian source populations.

## 2 Materials and Methods

### 2.1 Samples

Twenty *Ae. albopictus* were collected at Australian borders between April 4^th^, 2016, and December 17^th^, 2018. These mosquitoes, hereafter called ‘incursives’, were intercepted at passenger and freight terminals at airports and seaports in Australia, with most collected at marine terminals. Table 1 contains information for each of these 20 incursives. Overall, 2 *Ae. albopictus* (representing 2 independent detections) were collected at airports, and 18 *Ae. albopictus* (representing 10 independent detections) were collected at seaports. Incursives were collected at adult and immature life stages using several methods: adults were collected with BG-Sentinel traps (Biogents AG, Regensburg, Germany), BG-Gravid Aedes traps (Pacific Biologics, Scarborough, Queensland, Australia) or with aspirators; larvae and pupae were collected in tyre traps, ovitraps or through inspection of goods. Incursives were identified to species using morphology which was later confirmed in our genomic analysis.

**Table 1.** Detection and assignment results for the 9 well-assigned and 11 poorly-assigned *Aedes albopictus* incursives. Well-assigned incursives are in regular typeface, poorly-assigned incursives are in italics.

| Incursive | Port type | City of detection | Date of collection | Initial assignment | Final assignment | Posterior probability | Relative probability |
| --- | --- | --- | --- | --- | --- | --- | --- |
| AP01 | Airport | Sydney | 2016-04-30 | East Asia | China | 0.59 | 3.15 |
| AP02 | Airport | Melbourne | 2018-12-17 | East Asia | China | 0.77 | 5.50 |
| SP04 | Seaport | Darwin | 2017-06-05 | Southeast Asia | Singapore | 0.92 | 19.27 |
| SP05 | Seaport | Darwin | 2017-06-06 | Southeast Asia | Singapore | 0.91 | 11.43 |
| SP06 | Seaport | Brisbane | 2017-07-18 | Japan | Japan | 0.72 | 3.54 |
| SP09 | Seaport | Brisbane | 2018-03-26 | East Asia | Taiwan | 0.66 | 3.02 |
| SP15 | Seaport | Brisbane | 2018-10-24 | East Asia | China | 0.72 | 3.54 |
| SP16 | Seaport | Brisbane | 2018-10-24 | East Asia | China | 0.66 | 4.18 |
| SP18 | Seaport | Brisbane | 2018-10-26 | Japan | Japan | 0.82 | 6.09 |
| *SP01* | *Seaport* | *Brisbane* | *2016-12-23* | *East Asia* | *n/a* | *0.62* | *2.35* |
| *SP02* | *Seaport* | *Brisbane* | *2016-12-23* | *East Asia* | *n/a* | *0.51* | *2.37* |
| *SP03* | *Seaport* | *Brisbane* | *2016-12-23* | *East Asia* | *n/a* | *0.49* | *1.45* |
| *SP07* | *Seaport* | *Brisbane* | *2017-07-18* | *Japan* | *n/a* | *0.63* | *2.49* |
| *SP08* | *Seaport* | *Brisbane* | *2017-07-18* | *East Asia* | *n/a* | *0.48* | *1.73* |
| *SP10* | *Seaport* | *Brisbane* | *2018-03-29* | *East Asia* | *n/a* | *0.59* | *2.30* |
| *SP11* | *Seaport* | *Brisbane* | *2018-05-21* | *Japan* | *n/a* | *0.50* | *1.32* |
| *SP12* | *Seaport* | *Brisbane* | *2018-05-21* | *Japan* | *n/a* | *0.50* | *1.42* |
| *SP13* | *Seaport* | *Brisbane* | *2018-05-28* | *Japan* | *n/a* | *0.52* | *1.56* |
| *SP14* | *Seaport* | *Brisbane* | *2018-05-28* | *East Asia* | *n/a* | *0.64* | *2.83* |
| *SP17* | *Seaport* | *Brisbane* | *2018-10-24* | *East Asia* | *n/a* | *0.53* | *2.48* |

For the panel of reference genotypes, we used 241 *Ae. albopictus* collected from 18 locations in the Indo-Pacific region (Fig 1), with details listed in Supporting Information Appendix S1. These 18 locations were: China (Guangzhou), Christmas Island, Fiji, Japan (Matsuyama), Malaysia (Peninsula Malaysia), Mauritius, Philippines (Manila), Singapore, Sri Lanka (Colombo), Taiwan (Kaohsiung City), Thailand (Bangkok and Chiang Mai), Timor-Leste, Torres Strait Islands, Vanuatu, Vietnam (Ho Chi Minh City), and three locations in Indonesia: Bali, Bandung, and Jakarta. We considered different countries to represent different locations and aggregated mosquitoes accordingly, except in the case of Indonesia, where we treated samples from Bali, Bandung, and Jakarta as different locations. This was due to the proximity of Indonesia to Australia and the previous finding that most *Ae. aegypti* incursions into Australia were from Bali.^8^

**Fig 1.**
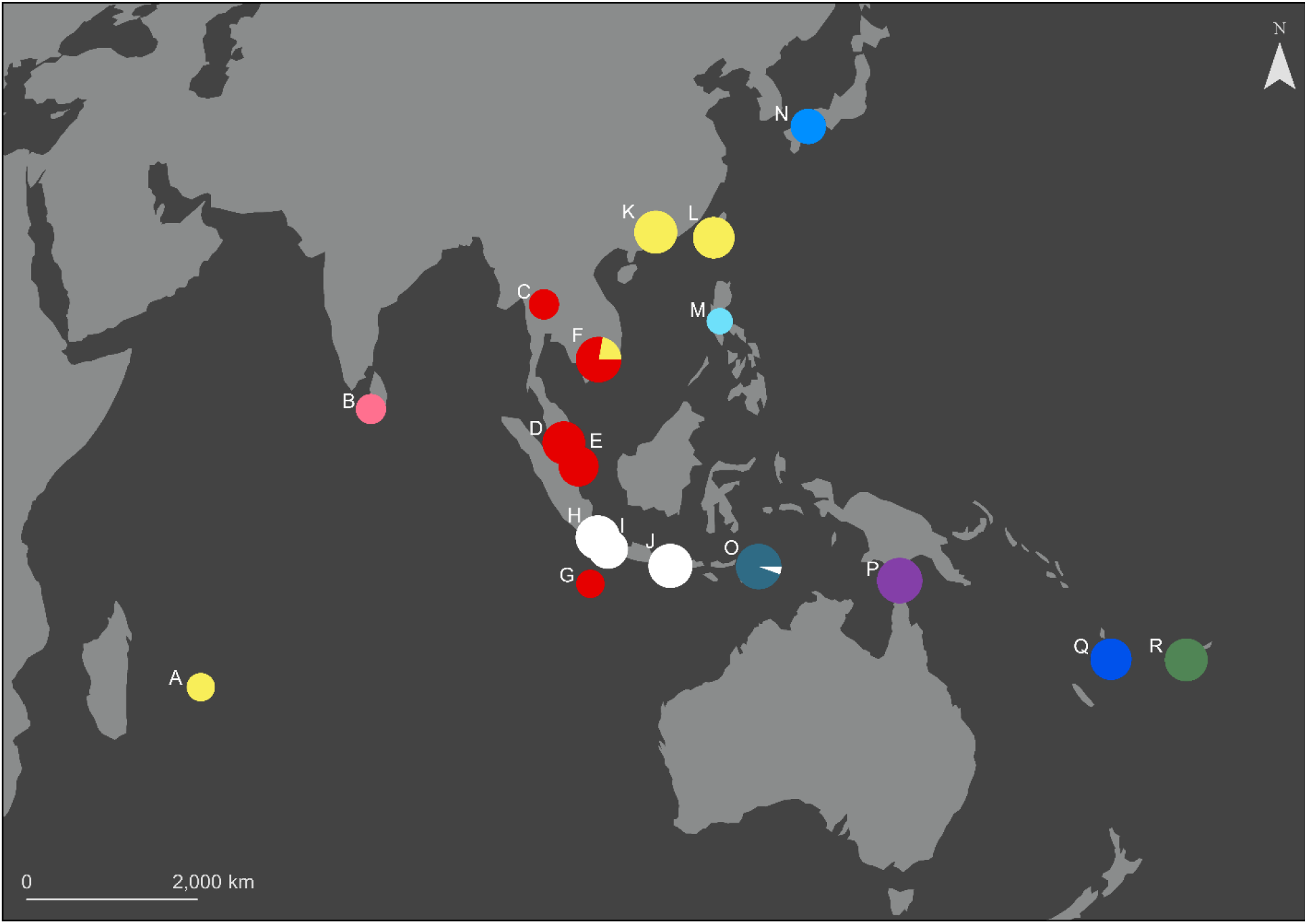
Reference genotypes of *Aedes albopictus*. Pie charts are sized by the number of reference genotypes available, and colours denote clustering established during the initial discriminant analysis of principal components (DAPC; see Results). Populations are: A) Mauritius; B) Sri Lanka; C) Thailand; D) Malaysia; E) Singapore; F) Vietnam; G) Christmas Island; H) Jakarta, Indonesia; I) Bandung, Indonesia; J) Bali, Indonesia; K) China; L) Taiwan; M) Philippines; N) Japan; O) Timor-Leste; P) Torres Strait Islands; Q) Vanuatu; and R) Fiji.

### 2.2 Genotyping

DNA was extracted from incursive and reference mosquitoes using either Qiagen DNeasy Blood & Tissue Kits (Qiagen, Hilden, Germany) or Roche High Pure™ PCR Template Preparation Kits (Roche Molecular Systems, Inc., Pleasanton, CA, USA), each with an RNase A treatment step.

We constructed double digest restriction-site associated DNA (ddRAD) libraries following the ddRAD protocol for *Ae. albopictus* adapted in Schmidt et al.^29^ from the protocol of Rašić, Filipović, Weeks, & Hoffmann.^30^ We performed initial digestions of 50 – 200 ng of genomic DNA, using 10 units each of MluCI and NlaIII restriction enzymes (New England Biolabs, Beverly MA, USA), NEB CutSmart buffer, and water. Digestions were run for 3 hours at 37 °C with no heat kill step, and the products were cleaned with paramagnetic beads. These were ligated to modified Illumina P1 and P2 adapters overnight at 16 °C with 1,000 units of T4 ligase (New England Biolabs, Beverly, MA, USA), followed by a 10-minute heat-deactivation step at 65 °C. We performed size selection with a Pippin-Prep 2% gel cassette (Sage Sciences, Beverly, MA) to retain DNA fragments of 350 – 450 bp.

Final libraries were amplified by PCR, using 1 μL of size-selected DNA, 5 μL of Phusion High Fidelity 2× Master mix (New England Biolabs, Beverly MA, USA) and 2 μL of 10 μM standard Illumina P1 and P2 primers. These were run for 12 PCR cycles, then cleaned and concentrated using 0.8× paramagnetic beads. Each ddRAD library contained between 30 and 65 mosquitoes, and each was sequenced on a single sequencing lane using 100 bp chemistry. Libraries were sequenced paired-end at either the Australian Genome Research Facility (AGRF, Melbourne, Australia), the University of Melbourne (Pathology), or GeneWiz, Inc (Suzhou, China) on either a HiSeq 2500 or a HiSeq 4000 (Illumina, California, USA).

Raw sequence reads were processed using the Stacks 2.0 REF_MAP pipeline.^31^ Reads were de-multiplexed and trimmed to 80 bp length, and low-quality reads were discarded. Bowtie 2.3.4.3^32^ was used to align reads to the *Aedes albopictus* reference genome (GenBank assembly accession: GCA_006496715.1) using end-to-end alignment and the --sensitive preset options. The 20 incursive individuals and 241 reference individuals were built into a Stacks loci catalog and the Stacks program POPULATIONS was used to call SNPs and export genotypes as a VCF file. Only SNPs called in 75% of individuals from each reference population and in 75% of incursives were retained (−p 19 −r 0.75). We used VCFTOOLS^33^ to ensure no first-order relatives were included in the reference populations (--relatedness2; only including individuals with kinship scores < 0.177).

### 2.3 Population assignment

We used two R packages for population assignment: ‘adegenet v2.1.2’,^34^ for cluster detection with discriminant analysis of principal components (DAPC); and ‘assignPOP v1.1.9’,^35^ for Monte Carlo assignment tests using a support vector machine predictive model. Our methodology was adapted from that of Schmidt et al.^8^ as follows. First, we performed DAPC on the whole dataset to partition the reference and incursive genotypes into broad genetic clusters, which provided an initial, rough assignment for each incursive. We then analysed each genetic cluster in isolation, and used assignPOP to generate assignment probabilities for each incursive to each of the reference populations within the genetic cluster.

For the initial analysis with DAPC, we used all 261 genotypes. These were filtered with POPULATIONS to retain SNPs scored in 75% of individuals from each reference population and in 75% of incursives. The dataset was then filtered with VCFTOOLS using the following settings: SNPs must be biallelic; have a minor allele count ≥ 2;^36^ have a minimum read depth of 3 and a maximum of 45 ((mean depth) + 3√(mean depth));^37^ and must be called in at least 90% of individuals. We then used BEAGLE 4.1^38^ to impute missing genotypes, with 10 iterations.

We used the *find.clusters* function to detect genetic clusters, with 10^9^ iterations. We ran cluster detection separately for each incursive, so that each incursive genotype was compared with all reference genotypes and none of the other incursive genotypes. We used *find.clusters* to partition the genotypes into a number of clusters (*K*) ranging from 1 to 20, and considered the *K* with the lowest Akaike Information Criterion (AIC) as the best partitioning. Following this partitioning, the dataset was divided into *K* clusters, each containing a set of reference genotypes and between 0 and 20 incursives.

The genetic clusters that contained at least one incursive were then analysed independently. New VCF files for each cluster were exported following the same filtering steps described above. We used the assignPOP function *assign.X* to generate posterior probabilities of assignment for incursives to the reference populations within their genetic cluster, using a support vector machine predictive model. Following Schmidt et al.,^8^ we applied these posterior probabilities to calculate ‘relative probabilities’ of assignment, defined as the probability of assignment to the most likely population divided by the probability of assignment to the second most likely population. We considered incursives assigned with posterior probability > 0.5 and relative probability > 3 to be well-assigned.

We performed final DAPCs using these assignments, with genotypes grouped by population of origin (for reference genotypes) or population of assignment (for well-assigned incursives). We omitted poorly-assigned incursives from these DAPCs. For each DAPC, we used the function *xvalDapc* with 100 repetitions to determine an optimal number of principal components to use, based on lowest mean squared error. DAPCs for each genetic cluster were plotted to investigate relationships between reference and incursive genotypes graphically.

We used the assignment results to compare incursion pathways of *Ae. albopictus* with those of *Ae. aegypti* established in Schmidt et al..^8^ For each species, we plotted the incursion routes (source population to interception location) into Australia of all well-assigned incursives.

## 3 Results

After filtering, we retained 21,931 SNPs for the initial analysis with DAPC. Lowest AIC was observed at *K* = 10. Accordingly, the reference genotypes were partitioned into 10 genetic clusters (Fig 1), and the incursives were assigned to these 10 clusters one by one. Only three of the clusters were assigned incursives. Twelve of these were assigned to an “East Asian” cluster containing China, Mauritius, Taiwan, and four of the Vietnamese genotypes (Fig 1, yellow). Two were assigned to a “Southeast Asian” cluster containing *Ae. albopictus* from Christmas Island, Malaysia, Singapore, Thailand, and the remaining 14 Vietnamese genotypes (Fig 1, red). The other six incursives were assigned to a “Japanese” cluster containing the Japanese genotypes.

More precise assignments were obtained by analysing each cluster separately, using assignPOP to produce posterior and relative probabilities of assignment for each incursive. Although the Vietnamese genotypes were split between the East Asian and Southeast Asian clusters, we included all Vietnamese genotypes in analyses of each cluster. Also, as multiple populations were needed for assignment tests, we included the Chinese and Taiwanese genotypes in analyses of the Japanese cluster, due to their geographic proximity with Japan. After filtering the East Asian cluster, there remained 14,149 SNPs for analysis. Analysis of the Southeast Asian cluster was with 12,346 SNPs, and the Japanese cluster was analysed with 26,980 SNPs.

Five of the 12 incursives initially assigned to the East Asian cluster were considered well-assigned (posterior probability > 0.5 and relative probability > 3) (Table 1). These incursives were all assigned to either China (4) or Taiwan (1), with none assigned to Mauritius or Vietnam. Graphical investigation of assignments using DAPC (Fig 2a) showed that two of the less-confidently assigned incursives (i.e. those displayed as smaller white squares) had some ambiguity of assignment between China (red circles) and Taiwan (blue circles). This suggests that the genetic similarity of these two populations may make precise assignments between them difficult.

**Fig 2.**
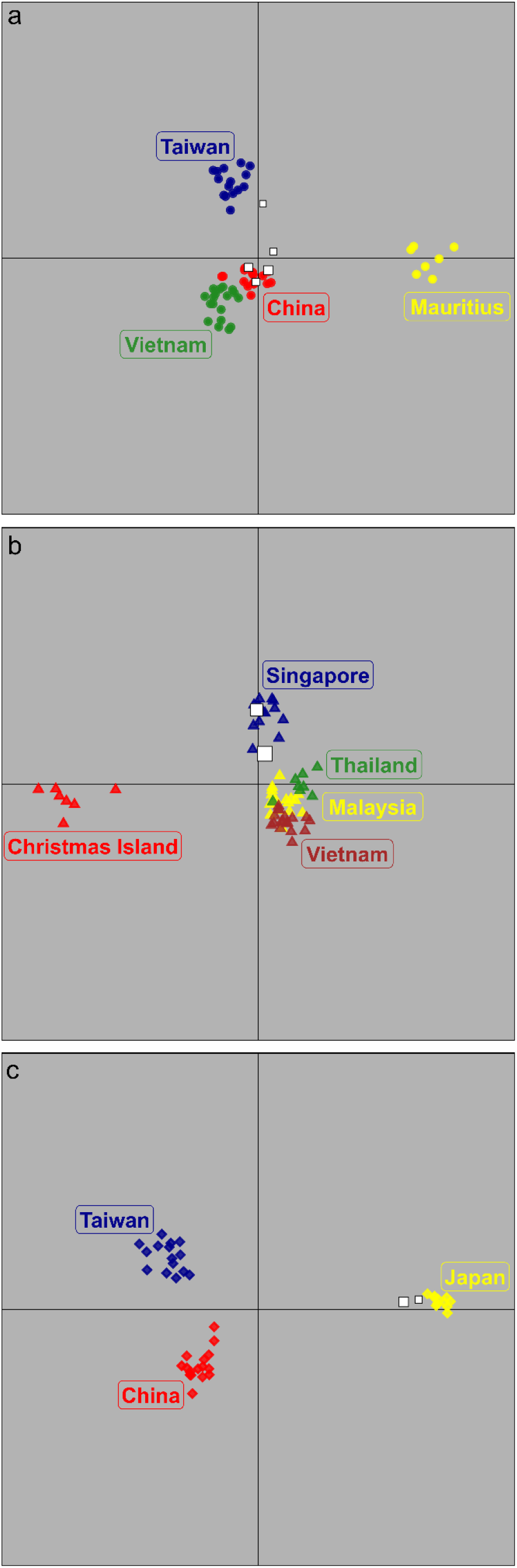
Discriminant analysis of principal components (DAPC) of the three clusters. White squares depict well-assigned incursive genotypes, which are sized by a logarithmic function of their relative probability of assignment. (a) East Asian cluster, horizontal axis describes 61.6% of total variation, vertical axis describes 26.5% of total variation; (b) Southeast Asian cluster, horizontal axis describes 60.3% of total variation, vertical axis describes 21.4% of total variation; (c) Japanese cluster, horizontal axis describes 88.3% of total variation, vertical axis describes 11.7% of total variation.

The two incursives initially assigned to the Southeast Asian cluster were both assigned very confidently to Singapore (both posterior probabilities > 0.9, relative probabilities > 11) (Table 1). DAPC of this cluster (Fig 2b) indicated that the confidence of these assignments was likely due in part to the clear differentiation of the Singaporean genotypes (blue triangles) from those from Malaysia (yellow triangles), Thailand (green triangles), and Vietnam (brown triangles), which clustered together. Of the six incursives initially assigned to the Japanese cluster, two were well-assigned (Table 1). Graphical investigation of these assignments indicated that these incursives were unlikely to have come from either China or Taiwan (Fig 2c).

*Aedes albopictus* incursion routes into Australia were clearly different from those of *Ae. aegypti* (Fig 3, Table 2). *Aedes aegypti* incursives from international sources were assigned to Brazil (8%, not pictured in Fig 3) and three locations in the Indo-Pacific: Bali (75%), Malaysia (16%), and Thailand (1%). Although reference genotypes from these three Indo-Pacific locations were used in this study, no *Ae. albopictus* incursives were assigned to these populations. Likewise, no *Ae. aegypti* were assigned to either Singapore or Taiwan, despite available reference genotypes from these populations. Overall, most *Ae. albopictus* incursion pathways were from East Asian sources into ports in eastern Australia, particularly Brisbane, and all 11 poorly-assigned incursives were collected at Brisbane. Most *Ae. aegypti* incursives were from Southeast Asia, and were detected at ports throughout Australia (but not Brisbane).

**Fig 3.**
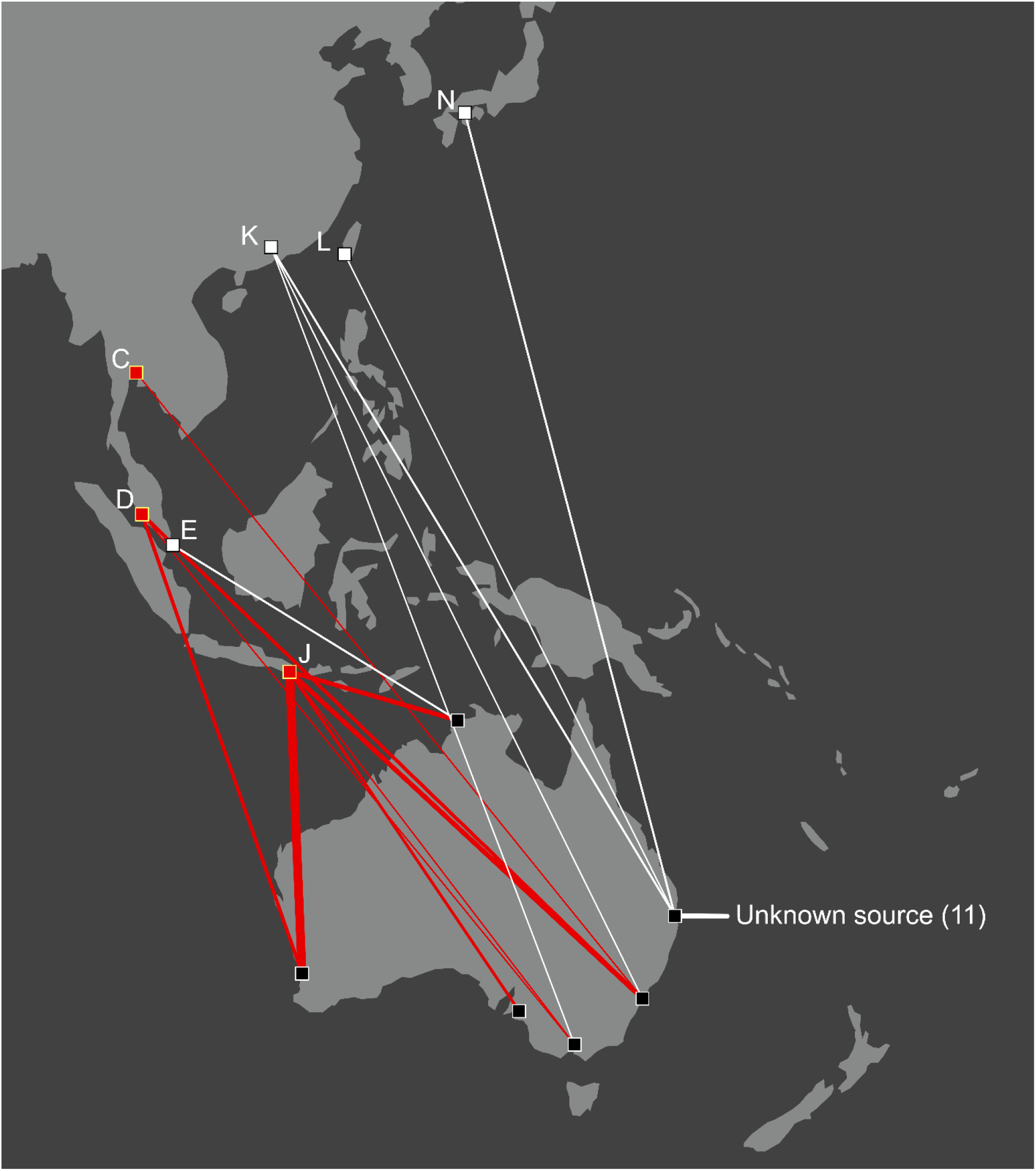
Indo-Pacific incursion pathways of *Aedes albopictus* (white) and *Aedes aegypti* (red). Line thickness follows a logarithmic function of the number of incursives transported along that pathway. Assignments for *Aedes aegypti* incursions are from Schmidt et al..^8^ Not shown are seven *Aedes aegypti* incursives, transported into Perth, Australia, from a putative origin in the Americas.

**Table 2.** Proportions of incursives assigned to each possible source population, for *Aedes albopictus* and *Aedes aegypti*. Assignments for *Aedes aegypti* incursions are from Schmidt et al..^8^

|  | *Aedes albopictus* | *Aedes aegypti* |
| --- | --- | --- |
| China | 0.444 | n/a |
| Singapore | 0.222 | 0 |
| Taiwan | 0.111 | 0 |
| Japan | 0.222 | n/a |
| Bali | 0 | 0.747 |
| Malaysia | 0 | 0.157 |
| Brazil | n/a | 0.084 |
| Thailand | 0 | 0.012 |
| Fiji | 0 | 0 |
| Sri Lanka | 0 | 0 |
| Vanuatu | 0 | 0 |
| Vietnam | 0 | 0 |
| Bandung | 0 | n/a |
| Christmas Island | 0 | n/a |
| Jakarta | 0 | n/a |
| Mauritius | 0 | n/a |
| Philippines | 0 | n/a |
| Timor-Leste | 0 | n/a |
| Torres Strait Islands | 0 | n/a |
| Kiribati | n/a | 0 |
| New Caledonia | n/a | 0 |
| Yogyakarta | n/a | 0 |

## 4 Discussion

This study has identified incursion pathways of *Ae. albopictus* mosquitoes into Australia, and shown that these pathways are strikingly different from those of its close relative, *Ae. aegypti*. We identified four locations from which intercepted *Ae. albopictus* were transported to Australia: three in East Asia (China, Taiwan, Japan) and one in Southeast Asia (Singapore). No Australian *Ae. aegypti* incursions have been linked to any of these locations, despite the presence of *Ae. aegypti* in both Taiwan and Singapore.^8^ Likewise, no *Ae. albopictus* in this study were found to have come from either Indonesia or Malaysia, the most common source populations of *Ae. aegypti*. These results show how even congeneric taxa with similar capacity for active dispersal can differ greatly in the dispersal pathways they take. Given that both *Ae. albopictus* and *Ae. aegypti* represent dangerous pest disease vectors and quarantine risks, the work highlights the importance of evaluating incursion pathways of pest species individually rather than by assuming that related species will have similar incursion pathways. For Australia and other countries yet to be invaded by *Ae. albopictus*, our findings should help current efforts to prevent further spread of this species. Additionally, with both species carrying insecticide resistance mutations,^39,40^ the identification of incursion pathways can help to reduce the risk of resistance alleles spreading into resident populations. For instance, in *Ae. aegypti* resistance alleles have spread between geographically distant populations within the Indo-Pacific region,^5^ highlighting the risk involved in genetic incursions as well as species incursions into areas where resistance alleles are absent.^41^

Using the composite methodology of DAPC and assignment tests, 9 of the 20 *Ae. albopictus* were well-assigned (posterior probability > 0.5 and relative probability > 3). This proportion (45%) is lower than the proportion of well-assigned *Ae. aegypti* in Schmidt et al.^8^ (76%), which is likely in part due to this study using a more conservative cut-off for assignment than used in Schmidt et al.^8^ (posterior probability > 0.5 and relative probability > 2). The methodology here proved sufficient for confidently assigning among the genetically and geographically close populations of Singapore and Malaysia (FST’ = 0.0251^20^), with the assignments to Singapore having the highest posterior and relative probabilities of all incursives (Table 1). Likewise, confident assignments were made to Japan, the source of *Ae. albopictus* incursions into Townsville, Australia, in 2012.^14^ One of these incursives (SP18) was detected in a shipment of used tyres originally from Japan, providing a clear concordance between genetic inferences and observations at point of detection.

Among the poorly-assigned incursives were three *Ae. albopictus* (SP01-03) intercepted on a used power boat that was brought to Brisbane from Florida, USA, and four *Ae. albopictus* (SP11-14) found in a shipment of used tyres from the USA that transhipped through Colombia. These were initially assigned to the East Asian cluster with low confidence (Table 1). In comparison, two of three *Ae. albopictus* (SP15-17) detected in fish bins from Hong Kong and one incursive (AP01) detected on cut-flowers from China were well-assigned to China, despite the high genetic similarity of Chinese and Taiwanese reference populations (FST’ = 0.0197^20^). Considering these results, the low confidence in specific assignments for incursives SP01-03 and SP11-14 may reflect the absence of reference genotypes from the Americas. *Aedes albopictus* in North America have genetic similarities with East Asian populations,^25^ and North American incursions of *Ae. albopictus* have been linked to goods transported from South China.^42^ As our databank of reference genotypes grows and we are able to add populations from the Americas, we may find some of these incursives are confidently assigned to North America.

Despite the absence of American reference genotypes, the methodology used in this study is sufficient to show clear differences in incursion pathways between this species and *Ae. aegypti* (Fig 5). In contrast to *Ae. aegypti*, for which most incursives are transported to Australia from popular tourism locations like Bali, the *Ae. albopictus* incursions investigated in this study came from locations that, while popular as tourist destinations, are also major trading partners of Australia (https://dfat.gov.au/trade/resources/trade-at-a-glance/). We estimate that 90% of *Ae. albopictus* incursives came to Australia by marine routes, compared with only 5% of *Ae. aegypti*,^8^ though the sample size for *Ae. albopictus* is currently small at 20 individuals and these patterns may change as more incursives are detected. Other observations linking *Ae. albopictus* with marine incursions and *Ae. aegypti* with aerial incursions have been noted at New Zealand ports.^17^ Considering the broad sympatry of these species throughout the Indo-Pacific,^43^ these findings raise questions about why marine incursions are relatively rare for *Ae. aegypti* and why aerial incursions are relatively rare for *Ae. albopictus*.

The low incidence of marine incursions for *Ae. aegypti* may reflect a lack of diapause in this species.^24^ By undergoing diapause, *Ae. albopictus* may be able to withstand longer transportation times and colder minimum temperatures than *Ae. aegypti,*^21^ which may increase its likelihood of surviving marine transportation to Australia. Thus even if a vessel is transporting similar numbers of *Ae. albopictus* and *Ae. aegypti* at departure, only *Ae. albopictus* may survive long marine voyages. This is concordant with the patterns of clustering observed in this study showing genetic similarities between East Asian and Mauritian *Ae. albopictus* populations (Fig 1). Given the very high frequency of international shipping past Mauritius,^20^ similarity of these locations likely reflects recent gene flow associated with marine transportation. Similar patterns of genetic clustering among distant populations has not been observed in *Ae. aegypti*.

A tentative explanation for the paucity of aerial incursions of *Ae. albopictus* into Australia and New Zealand may relate to aircraft disinsection^44^ and insecticide resistance. All international flights into Australia and New Zealand undergo disinsection with synthetic pyrethroids,^45^ a class of insecticide to which both species have developed some resistance via mutations at the *Vssc* gene.^39,40^ All *Ae. aegypti* incursives into Australia and New Zealand carry these resistance mutations;^8^ in Bali, the most common source population for *Ae. aegypti* incursions, every mosquito was homozygous for a pair of resistance mutations conferring strong resistance to Type I and Type II synthetic pyrethroids. Although homologous resistance mutations putatively conferring Type I pyrethroid resistance have been recorded in *Ae. albopictus*,^46^ there have been no reports of mutations for Type II resistance and it is unclear how widely distributed the homologous mutations are in the Indo-Pacific region. If *Ae. albopictus* from locations such as Bali remain susceptible to certain types of pyrethroids, aircraft disinsection may be more effective at killing *Ae. albopictus* than *Ae. aegypti*. A comprehensive geographic investigation of resistance mutations in *Ae. albopictus* will be required to evaluate this hypothesis, and to test whether disinsection is more effective at stopping incursions of this species than *Ae. aegypti*. Other factors potentially influencing the relative frequency of aerial and marine incursions include any differences in relative abundance of *Ae. albopictus* and *Ae. aegypti* around airports and seaports and any differences in behaviour that affect the likelihood of mosquitoes boarding and travelling on vessels.

## Conclusions

Understanding pest incursion pathways is critical for preventing new invasions and stopping the transfer of alleles that reduce the efficacy of control methods. This study has traced incursion pathways of *Ae. albopictus* into mainland Australia, where no successful invasions have been observed. Most strikingly, we found that the source populations of *Ae. albopictus* incursions were all different to those of the closely-related mosquito *Ae. aegypti*, despite broad sympatry of these species in the Indo-Pacific region. Incursions of *Ae. albopictus* came mostly along marine routes, from key trading locations, while *Ae. aegypti* mostly came by aerial routes from tourism hotspots. As with our previous research on *Ae.* aegypti,^8^ our new results for *Ae. albopictus* highlight the value of genomics for deciphering otherwise cryptic incursion pathways. Of particular importance are the findings from the comparative analysis of both datasets. Based on the different pathways of the two species, we strongly recommend against assuming that closely-related taxa will share common incursion pathways, and propose that incursion pathways of important taxa should be investigated individually. Management plans for *Ae. albopictus* and *Ae. aegypti* should incorporate knowledge of how insecticide resistance directs and modifies incursion pathways, and how it is transmitted along incursion pathways into areas where resistance alleles are absent.

## Acknowledgements

We thank Qiong Yang and Katie Robinson for molecular lab assistance. We thank Esther Anderson, Ashley Callahan, Gerhard Ehlers, Joe Davis and Odwell Muzari for sample collection. We thank the Department of Agriculture, Water and the Environment for providing funding for this work. AAH was supported by Program and Fellowship grants from the National Health and Medical Research Council (NHMRC). TLS and AAH were also supported by the Wellcome Trust UK.

## References

1. Gubler D and Kuno G, Dengue and dengue hemorrhagic fever: its history and resurgence as a global public health problem, in Dengue and Dengue Hemorrhagic Fever. CAB international, London, pp. 1–22 (1997).

2. Walsh JR, Carpenter SR and Van Der Zanden MJ, Invasive species triggers a massive loss of ecosystem services through a trophic cascade. Proc Natl Acad Sci 38: 4081–4085 (2016).

3. Paini DR, Sheppard AW, Cook DC, De Barro PJ, Worner SP and Thomas MB, Global threat to agriculture from invasive species. Proc Natl Acad Sci 113: 7575–7579 (2016).

4. Gao, Y and Reitz, S, Emerging Themes in Our Understanding of Species Displacements. Annu Rev Entomol 62: 165–183 (2017).

5. Endersby-Harshman NM, Schmidt TL, Chung J, Rooyen A van, Weeks AR and Hoffmann AA, Heterogeneous genetic invasions of three insecticide resistance mutations in IndoPacific populations of *Aedes aegypti* (L.). Mol Ecol 29: 1628–1641 (2020).

6. Levine JM and D’Antonio CM, Forecasting biological invasions with increasing international trade. Conserv Biol 17: 322–326 (2003).

7. Westphal MI, Browne M, MacKinnon K and Noble I, The link between international trade and the global distribution of invasive alien species. Biol Invasions 10: 391–398 (2008).

8. Schmidt TL, van Rooyen AR, Chung J, Endersby‐Harshman NM, Griffin PC, Sly A, Hoffmann AA and Weeks, AR, Tracking genetic invasions: Genome‐wide single nucleotide polymorphisms reveal the source of pyrethroid‐resistant *Aedes aegypti* (yellow fever mosquito) incursions at international ports. Evol Appl 12:1136–1146 (2019).

9. Hulme PE, Trade, transport and trouble: managing invasive species pathways in an era of globalization. J Appl Ecol 46: 10–18 (2009).

10. Caley P, Ingram R and De Barro P. Entry of exotic insects into Australia: Does border interception count match incursion risk? Biol Invasions 17: 1087–1094 (2015).

11. Robinson A, Burgman MA and Cannon R, Allocating surveillance resources to reduce ecological invasions: maximizing detections and information about the threat. Ecol Appl 21: 1410–1417 (2011).

12. Mehta S V., Haight RG, Homans FR, Polasky S and Venette RC, Optimal detection and control strategies for invasive species management. Ecol Econ 61: 237–245 (2007).

13. Hawley WA, The biology of *Aedes albopictus*. J Am Mosq Control Assoc Suppl 1: 1–39 (1988).

14. van den Hurk AF, Nicholson J, Beebe NW, Davis J, Muzari OM, Russell RC, Devine GJ and Ritchie SA, Ten years of the Tiger: *Aedes albopictus* presence in Australia since its discovery in the Torres Strait in 2005. One Heal 2: 19–24 (2016).

15. Kraemer MUG, Reiner RC, Brady OJ, Messina JP, Gilbert M, Pigott DM, Yi D, Johnson K, Earl L, Marczak LB, Shirude S, Weaver ND, Bisanzio D, Perkins TA, Lai S, Lu X, Jones P, Coelho GE, Carvalho RG, Bortel WV, Marsboom C, Hendrickx G, Schaffner F, Moore CG, Nax HH, Bengtsson L, Wetter E, Tatem AJ, Brownstein JS, Smith DL, Lambrechts L, Cauchemez S, Linard C, Faria NR, Pybus OG, Scott TW, Liu Q, Yu H, Wint GRW, Hay SI and Golding N, Past and future spread of the arbovirus vectors *Aedes aegypti* and *Aedes albopictus*. Nat Microbiol 4: 854–863 (2019).

16. Ritchie SA, Moore P, Carruthers M, Williams C, Montgomery B, Foley P, Ahboo S, van den Hurk AF, Lindsay MD, Cooper B, Beebe N and Russell RC, Discovery of a Widespread Infestation of *Aedes albopictus* in the Torres Strait, Australia. J Am Mosq Control Assoc 22: 358–365 (2006).

17. Ammar SE, Mclntyre M, Swan T, Kasper J, Derraik JGB, Baker MG and Hales S, Intercepted Mosquitoes at New Zealand’s Ports of Entry, 2001 to 2018: Current Status and Future Concerns. Trop Med Infect Dis 4: 101–118 (2019).

18. Lounibos LP, Invasions by Insect Vectors of Human Disease. Annu Rev Entomol 47: 233–266 (2002).

19. Tatem AJ, Hay SI and Rogers DJ, Global traffic and disease vector dispersal. Proc Natl Acad Sci 103: 6242–6247 (2006).

20. SchmidtTL, Chung J, Honnen, A-C, Weeks AR and Hoffmann AA, Population genomics of two invasive mosquitoes (*Aedes aegypti* and *Aedes albopictus*) from the Indo-Pacific. bioRxiv DOI: 10.1101/2020.03.15.993055.

21. Urbanski JM, Benoit JB, Michaud MR, Denlinger DL and Armbruster P, The molecular physiology of increased egg desiccation resistance during diapause in the invasive mosquito, *Aedes albopictus*. Proc R Soc B Biol Sci 277: 2683–2692 (2010).

22. Medley KA, Westby KM and Jenkins DG, Rapid local adaptation to northern winters in the invasive Asian tiger mosquito *Aedes albopictus*: A moving target. J Appl Ecol 56: 2518–2527 (2019).

23. Diniz DFA, De Albuquerque CMR, Oliva LO, De Melo-Santos MAV and Ayres CFJ, Diapause and quiescence: Dormancy mechanisms that contribute to the geographical expansion of mosquitoes and their evolutionary success. Parasit Vectors 10: 310–322 (2017).

24. Mitchell C, Geographic spread of *Aedes albopictus* and potential for involvement in arbovirus cycles in the Mediterranean basin. J Vector Ecol 20: 44–58 (1995)

25. Kotsakiozi P, Richardson JB, Pichler V, Favia G, Martins AJ, Urbanelli S, Armbruster PA and Caccone A, Population genomics of the Asian tiger mosquito, *Aedes albopictus*: insights into the recent worldwide invasion. Ecol Evol 7: 10143–10157 (2017).

26. Manni M, Guglielmino CR, Scolari F, Vega-Rúa A, Failloux A-B, Somboon P, Lisa A, Savini G, Bonizzoni M, Gomulski LM, Malacrida AR and Gasperi G, Genetic evidence for a worldwide chaotic dispersion pattern of the arbovirus vector, *Aedes albopictus*. PLoS Negl Trop Dis 11: e0005332 (2017).

27. Sherpa S, Blum MGB, Capblancq T, Cumer T, Rioux D and Després L, Unravelling the invasion history of the Asian tiger mosquito in Europe. Mol Ecol, 28: 2360–2377 (2019).

28. Eskildsen GA, Rovira JR, Smith O, Miller MJ, Bennett KL, McMillan WO and Loaiza J, Maternal invasion history of *Aedes aegypti* and *Aedes albopictus* into the Isthmus of Panama: implications for the control of emergent viral disease agents. PLoS One 13: e0194874 (2018).

29. Schmidt TL, Rašić G, Zhang D, Zheng X, Xi Z and Hoffmann AA. Genome-wide SNPs reveal the drivers of gene flow in an urban population of the Asian Tiger Mosquito, *Aedes albopictus*. PLoS Negl Trop Dis 11: e0006009 (2017).

30. Rašić G, Filipović I, Weeks AR and Hoffmann AA. Genome-wide SNPs lead to strong signals of geographic structure and relatedness patterns in the major arbovirus vector, *Aedes aegypti*. BMC Genomics 15: 275–286 (2014).

31. Catchen J, Hohenlohe PA, Bassham S, Amores A and Cresko WA. Stacks: an analysis tool set for population genomics. Mol Ecol 22: 3124–3140 (2013).

32. Langmead B and Salzberg SL. Fast gapped-read alignment with Bowtie 2. Nat Methods 9: 357–359 (2012).

33. Danecek P, Auton A, Abecasis G, Albers CA, Banks E, DePristo MA, Handsaker RE, Lunter G, Marth GT, Sherry ST, McVean G, Durbin R and 1000 Genomes Project Analysis Group, The variant call format and VCFtools. Bioinformatics 27: 2156–2158 (2011).

34. Jombart T, adegenet: a R package for the multivariate analysis of genetic markers. Bioinformatics 24: 1403–1405 (2008).

35. Chen K-Y, Marschall EA, Sovic MG, Fries AC, Gibbs HL and Ludsin SA, *assignPOP*: An R package for population assignment using genetic, non-genetic, or integrated data in a machine-learning framework. Methods Ecol Evol 9: 439–446 (2018).

36. Linck E and Battey CJ, Minor allele frequency thresholds strongly affect population structure inference with genomic datasets. Mol Ecol Resour 19: 639–647 (2019).

37. Li H, Toward better understanding of artifacts in variant calling from high-coverage samples. Bioinformatics 30: 2843–2851 (2014).

38. Browning BL and Browning SR, Genotype Imputation with Millions of Reference Samples. Am J Hum Genet 98: 116–126 (2016).

39. Smith LB, Kasai S and Scott JG, Pyrethroid resistance in *Aedes aegypti* and *Aedes albopictus*: Important mosquito vectors of human diseases. Pestic Biochem Physiol 133: 1–12 (2016).

40. Auteri M, La Russa F, Blanda V and Torina A, Insecticide Resistance Associated with *kdr* Mutations in *Aedes albopictus*: An Update on Worldwide Evidences. Biomed Res Int 2018: 1–10 (2018).

41. Endersby-Harshman NM, Wuliandari JR, Harshman LG, Frohn V, Johnson BJ, Ritchie SA and Hoffmann AA, Pyrethroid Susceptibility Has Been Maintained in the Dengue Vector, *Aedes aegypti* (Diptera: Culicidae), in Queensland, Australia. J Med Entomol 54: 1649–1658 (2017).

42. Madon MB, Hazelrigg JE, Shaw MW, Kluh S and Mulla MS, Has *Aedes albopictus* established in California? J Am Mosq Control Assoc 19: 297–300 (2003).

43. Kraemer MUG, Sinka ME, Duda KA, Mylne AQN, Shearer FM, Barker CM, Moore CG, Carvalho RG, Coelho GE, Bortel WV, Hendrickx G, Schaffner F, Elyazar IRF, Teng H, Brady OJ, Messina JP, Pigott DM, Scott TW, Smith DL, Wint GRW, Golding N and Hay SI, The global distribution of the arbovirus vectors *Aedes aegypti* and *Ae. albopictus*. Elife 4: e08347 (2015).

44. IPCS, Aicraft disinsection insecticides (Environmental health criteria 243). European Commission and the Policy Research Programme, Geneva (2013).

45. DAWR/MPI, Schedule of aircraft disinsection procedures for flights into Australia and New Zealand. Department of Agriculture and Water Resources, Canberra, and Ministry for Primary Industries, Wellington (2016).

46. Xu J, Bonizzoni M, Zhong D, Zhou G, Cai S, Li Y, Wang X, Lo E, Lee R, Sheen R, Duan J, Yan G and Chen X, Multi-country Survey Revealed Prevalent and Novel F1534S Mutation in Voltage-Gated Sodium Channel (VGSC) Gene in *Aedes albopictus*. PLoS Negl Trop Dis 10: e0004696 (2016).

