## Supporting Information Appendix S1 for "Incursion pathways of the Asian tiger mosquito (*Aedes albopictus*) into Australia contrast sharply with those of the yellow fever mosquito (*Aedes aegypti*)"

| Label (Fig 1) | sample name | location(s) | country | year(s) collected | no. genotypes after filtering |
| --- | --- | --- | --- | --- | --- |
| 1 | Mauritius | Near Chamarel village | Mauritius | 2017 | 7 |
| 2 | Sri Lanka | Delgoda and Kiribathgoda, Colombo | Sri Lanka | 2017 | 8 |
| 3 | Thailand | Chiang Mai and Bangkok | Thailand | 2017 | 8 |
| 4 | Malaysia | Kuala Lumpur, Gombak, Kuantan and Johor Baru | Malaysia | 2015/2016 | 16 |
| 5 | Singapore | Near Bukit Timah Nature Reserve | Singapore | 2017 | 14 |
| 6 | Vietnam | Various locations in Ho Chi Minh City | Vietnam | 2017/2018 | 18 |
| 7 | Christmas Island | Near Christmas Island International Airport | Australia | 2017 | 7 |
| 8 | Jakarta | Various locations in Jakarta, Java | Indonesia | 2016 | 17 |
| 9 | Bandung | Various locations in Bandung, Java | Indonesia | 2017 | 14 |
| 10 | Bali | Various locations in Bali | Indonesia | 2016/2017 | 17 |
| 11 | China | 13 locations in Guangzhou | China | 2015 | 16 |
| 12 | Taiwan | Various locations in Southwest Taiwan | Republic of China | 2016 | 15 |
| 13 | Philippines | Manila, Quezon City and Makati City | Philippines | 2017 | 6 |
| 14 | Japan | Near Ehime University, Matsuyama | Japan | 2017 | 11 |
| 15 | Timor-Leste | Same | Timor-Leste | 2019 | 18 |
| 16 | Torres Strait Islands | 12 Inner and outer islands in the Torres Strait | Australia | 2018 | 18 |
| 17 | Vanuatu | Various locations in Efate | Vanuatu | 2018 | 15 |
| 18 | Fiji | Various locations in and near to Nadi | Fiji | 2018 | 16 |
